# Validation of the Social Responsiveness Scale (SRS) to screen for atypical social behaviors in juvenile macaques

**DOI:** 10.1101/2020.06.26.173161

**Authors:** Z. Kovacs Balint, J. Raper, V. Michopoulos, L.H. Howell, C. Gunter, J. Bachevalier, M.M. Sanchez

## Abstract

Primates form strong social bonds and depend on social relationships and networks that provide shared resources and protection critical for survival. Social deficits such as those present in autism spectrum disorder (ASD) and other psychiatric disorders hinder the individual’s functioning in communities. Given that early diagnosis and intervention can improve outcomes and trajectories of ASD, there is a great need for tools to identify early markers for screening/diagnosis, and for translational animal models to uncover biological mechanisms and develop treatments. One of the most widely used screening tools for ASD in children is the Social Responsiveness Scale (SRS), a quantitative measure used to identify individuals with atypical social behaviors. The SRS has been adapted for use in adult rhesus monkeys (*Macaca mulatta*) –a species very close to humans in terms of social behavior, brain anatomy/connectivity and development– but has not yet been validated or adapted for a necessary downward extension to younger ages matching those for ASD diagnosis in children. The goal of the present study was to adapt and validate the adult macaque SRS (mSRS) in juvenile macaques with age equivalent to 4-6 yr old human children. Expert primate coders modified the mSRS to adapt it to rate atypical social behaviors in juvenile macaques living in complex social groups at the Yerkes National Primate Research Center. Construct and face validity of this juvenile mSRS (jmSRS) was determined based on well-established and operationalized measures of social and non-social behaviors in this species using traditional behavioral observations. We found that the jmSRS identifies variability in social responsiveness of juvenile rhesus monkeys and shows strong construct/predictive validity, as well as sensitivity to detect atypical social behaviors in young male and female macaques across social status. Thus, the jmSRS provides a promising tool for translational research on macaque models of children social disorders.

## Introduction

Primates, both humans and nonhuman species, depend on social relationships that provide shared resources and protection critical for survival of the individual and the group. Social deficits present in neurodevelopmental disorders such as Autism Spectrum Disorder (ASD) and ADHD, and other psychiatric disorders (e.g. social anxiety, schizophrenia), can severely hinder the individual’s ability to function in a community. Social deficits impair daily functioning by altering social interactions, which can lead to social withdrawal and isolation (1). The prevalence of ASD, in particular, is estimated at 1 in 59 children in the USA (2). It is defined by impairments in social communication and interactions, in parallel to restricted interests and repetitive patterns of behavior (3). ASD is also associated with heightened anxiety and has a 4:1 male:female ratio in diagnoses (3, 4); however, recent research suggests that ASD diagnosis in females using “gold-standard” instruments may be significantly underestimating prevalence, for a variety of sex- and gender-based factors including “compensatory camouflaging” learned through socialization (5, American Psychiatric Association, 2013 #96). Early diagnosis and intervention is critical, given their positive effects in changing the trajectory of the disorder and improving outcomes (6, 7).

Identifying early biobehavioral markers of social deficits during development is critical for understanding ASD’s etiology and temporal unfolding, as well as for the development of early interventions and treatment. Currently, ASD can be diagnosed around 2 years of age (although the average age in the USA is 4 years old (2)), by observing deficits in social communication that develop as a child grows. The research and clinical standard to assess ASD and establish the diagnosis includes the Autism Diagnostic Interview–Revised (ADI-R), the Autism Diagnostic Observation Schedule (ADOS-2) and the Social Responsiveness Scale (SRS) (8-10). The SRS is a widely used questionnaire that quantitatively measures the continuum of both typical and atypical social behaviors that covary with ASD symptom severity and is completed by the caregivers (or a teacher) of the patient (8, 11). The items of the SRS measure a child’s ability to engage in reciprocal social interactions, and how deficits in communication and stereotypic/restricted interests and behaviors impair reciprocal social interactions in their naturalistic social environment (8).

The complex nature of ASD has made it hard to study its etiology and biological roots in human populations. Therefore, true progress in understanding neurobiological mechanisms underlying ASD will require examination of those processes in nonhuman primate (NHP) models with sophisticated social behaviors, phylogenetic closeness to humans and brain anatomy, connectivity, function and development that closely resemble that of our species. Rhesus monkeys (*Macaca mulatta*), in particular, have been widely used to understand the typical development of human social behavior, including the evolutionary context for Bowlby’s “Attachment Theory” (12-14). This species displays complex social behaviors, such as mother-offspring bonds, social play, reciprocal prosocial interactions, strong family alliances that preserve the status in the social hierarchy, as well as social awareness (15-18). The complexity of rhesus monkeys’ social behavior, their physiological, anatomical and genetic closeness to humans, and parallels in brain and social development (19-27) make them an ideal model organism to study typical and atypical development of human social function.

To leverage rhesus monkeys as a NHP animal model of translational value for ASD-related social deficits, it is important to first develop and validate screening tools across species and developmental stages. With this goal in mind, researchers at the Yerkes National Primate Research Center (YNPRC) first adapted one of the most widely used screening tools for ASD, the SRS (5, 8), for use in adult rhesus macaques (*macaque SRS:* mSRS (28)). The mSRS tested the construct of social responsiveness in adult macaques through an adaptation of the *“Chimpanzee SRS”*, which was, in turn, a cross-species adaptation of the human SRS to chimpanzees to measure social function (29). The *“Chimpanzee SRS”* detected variation in social behavior and its factor structure resembled that of the human SRS. The adult mSRS adapted from chimpanzees by Feczko et al (28) also identified variability in social responsiveness, was sensitive to detect atypical social behaviors, and showed a factor structure similar to the human and the chimpanzee SRS. An additional advantage of the mSRS is that, like the human SRS –which can be filled out by a caregiver or teacher without previous knowledge about the test or about the social responsiveness construct– the mSRS does not require previous knowledge/training on the social responsiveness construct or test. In addition, the mSRS takes a relatively small amount of time to complete by coders following traditional behavioral observations of social behavior of the animals.

However, the mSRS has not yet been validated or adapted for a necessary downward extension to younger ages matching those for ASD diagnosis in children. Since social deficits in ASD are rooted in early childhood, the development of a primate model to understand its etiology requires to first develop behavioral tools to identify typical and atypical social behaviors and underlying neurocircuitry in juvenile macaques. Therefore, the goal of the present study was to adapt and validate the adult mSRS in juvenile macaques with age equivalent to 4-6 yrs old human children. Developing an instrument that can translate to existing children measurements of social deficits, quickly completed following behavioral focal observations, and complementing the quantitative behavior collected using more established ethograms in our laboratory (30, 31) is highly needed. Based on our results, the juvenile mSRS (jmSRS) identifies variability in social responsiveness of juvenile rhesus monkeys and shows strong construct/predictive validity and sensitivity to detect atypical social behaviors in both male and female macaques across social status, providing a promising instrument for translational research on social disorders such as ASD.

## Methods

### Subjects and Housing

The subjects were 93 juvenile rhesus monkeys (*M. mulatta*) studied at approximately 1.5 yrs of age (Average: 17.32 ± 0.11months). All animals lived with their mothers and families in complex social groups at the YNPRC Field Station breeding colony (Lawrenceville, GA). Of the 93 juveniles studied, 44 were females (social rank distribution: 18 high, 14 middle, 12 low) and 49 were males (social rank distribution: 15 high, 18 middle and 13 low) from six different social groups consisting of 55-130 adult females with their adolescent, juvenile, and infant offspring, and 2-4 adult males. At this young age, social rank of the juvenile is based on their mothers’ matrilineal social rank. Matrilineal social rank was assessed from aggression and submission behaviors exhibited during dyadic agonistic interactions during group observations with high rank defined as the top 33% most dominant, low rank as the lowest 33%, and middle rank as those in between. The groups were housed in outdoor compounds with access to indoor climate-controlled housing areas. Animals were fed a standard commercial low-fat, high-fiber diet (Purina Mills International, LabDiets, St. Louis, MO) *ad libitum* in the morning and afternoon, supplemented each day with seasonal fruits or vegetables; water was freely available. Subjects were excluded from the study if they were from an unstable social group, or had clinical conditions at time of assignment. All procedures were approved by the Emory University Institutional Animal Care and Use Committee (IACUC), and were performed in accordance with the Animal Welfare Act and the U.S. Department of Health and Human Services “Guide for the Care and Use of Laboratory Animals” (32). The YNPRC is fully accredited by AAALAC International.

### Procedures

#### Behavioral data collection (Focal observations using rhesus ethogram)

Focal behavioral observations of the juveniles were collected by trained coders from observation towers situated over each social compound using a detailed ethogram adapted from a well-established rhesus monkey ethogram (33) and according to published methods (34-36). The ethogram captured frequencies and durations of affiliative (proximity, grooming, eye gaze, touch), agonistic (aggression, submission), anxiety-like (yawn, body shakes) and play behaviors exhibited by each subject towards other group members (social, solitary), as well as time spent alone and atypical behaviors similar to those seen in ASD (e.g. stereotypies) and other species- and age-specific behaviors. The full list of coded behaviors and their operational definitions are presented in **Table 1**. Data were collected using netbook computers with an in-house data acquisition software (WinObs60) by four trained observers with an inter-rater reliability of Cohen’s k=0.83. Subjects were identified by a distinctive dye-mark on their body, and each subject was focally observed for 30-min on four separate days (total of 2-hours of observation per subject) by one of the coders. Observations were collected between 0700 and 1200 hr, when animals were most active. All animals in the social group were kept outdoors during the observation sessions.

**Table 1.**
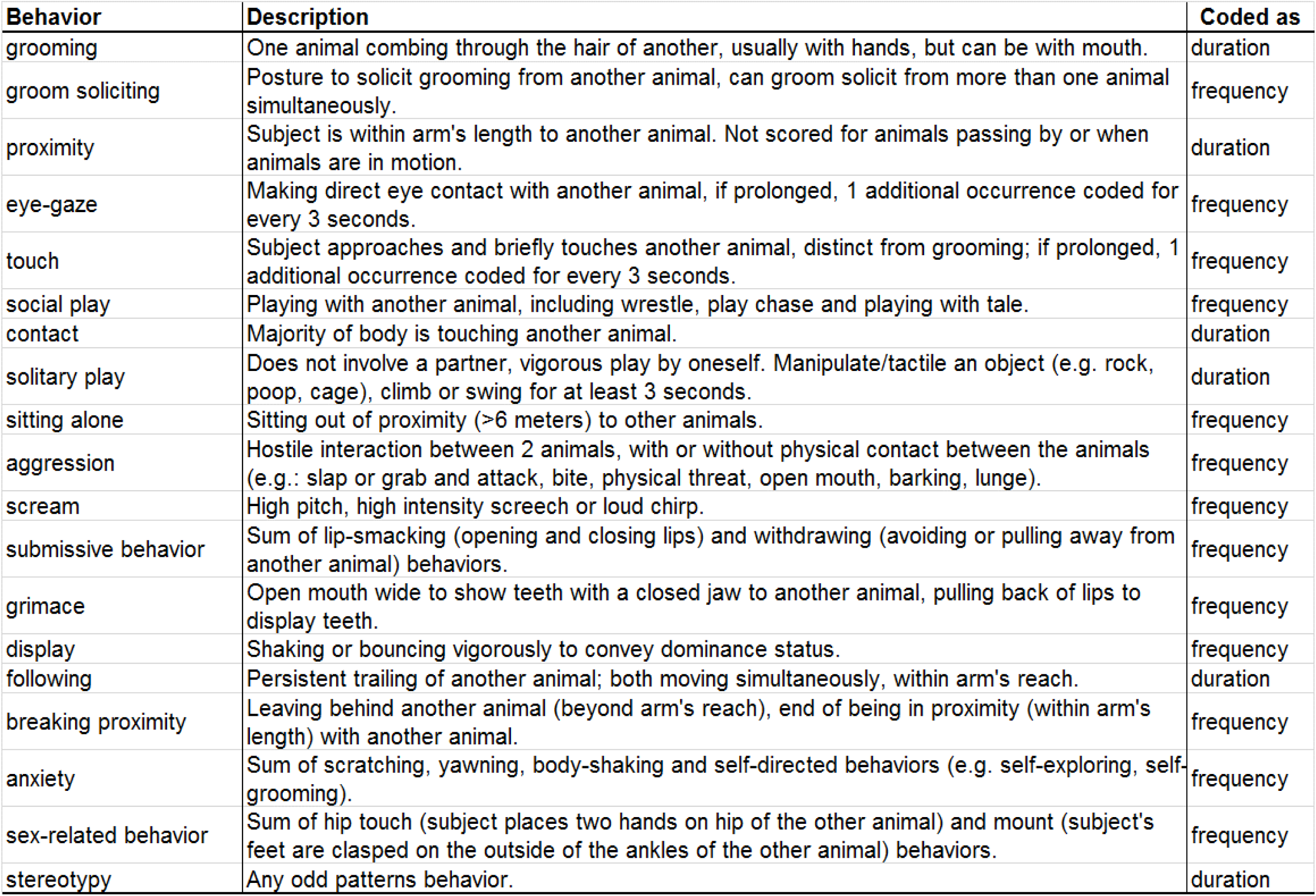
List of behaviors coded during the behavioral observations (ethogram). Approximately 2 hours of behavioral data was collected from each animal (on 4 different days) in their social compound.

| Behavior | Description | Coded as |
| --- | --- | --- |
| grooming | One animal combing through the hair of another, usually with hands, but can be with mouth. | duration |
| groom soliciting | Posture to solicit grooming from another animal, can groom solicit from more than one animal simultaneously. | frequency |
| proximity | Subject is within arm's length to another animal. Not scored for animals passing by or when animals are in motion. | duration |
| eye-gaze | Making direct eye contact with another animal, if prolonged, 1 additional occurrence coded for every 3 seconds. | frequency |
| touch | Subject approaches and briefly touches another animal, distinct from grooming; if prolonged, 1 additional occurrence coded for every 3 seconds. | frequency |
| social play | Playing with another animal, including wrestle, play chase and playing with tale. | frequency |
| contact | Majority of body is touching another animal. | duration |
| solitary play | Does not involve a partner, vigorous play by oneself. Manipulate/tactile an object (e.g. rock, poop, cage), climb or swing for at least 3 seconds. | duration |
| sitting alone | Sitting out of proximity (>6 meters) to other animals. | frequency |
| aggression | Hostile interaction between 2 animals, with or without physical contact between the animals (e.g.: slap or grab and attack, bite, physical threat, open mouth, barking, lunge). | frequency |
| scream | High pitch, high intensity screech or loud chirp. | frequency |
| submissive behavior | Sum of lip-smacking (opening and closing lips) and withdrawing (avoiding or pulling away from another animal) behaviors. | frequency |
| grimace | Open mouth wide to show teeth with a closed jaw to another animal, pulling back of lips to display teeth. | frequency |
| display | Shaking or bouncing vigorously to convey dominance status. | frequency |
| following | Persistent trailing of another animal; both moving simultaneously, within arm's reach. | duration |
| breaking proximity | Leaving behind another animal (beyond arm's reach), end of being in proximity (within arm's length) with another animal. | frequency |
| anxiety | Sum of scratching, yawning, body-shaking and self-directed behaviors (e.g. self-exploring, self-grooming). | frequency |
| sex-related behavior | Sum of hip touch (subject places two hands on hip of the other animal) and mount (subject's feet are clasped on the outside of the ankles of the other animal) behaviors. | frequency |
| stereotypy | Any odd patterns behavior. | duration |

#### Juvenile macaque social responsiveness scale (jmSRS)

A 14-item global rating instrument of typical and atypical social behaviors was used. This instrument was adapted from the adult rhesus macaque SRS described above (mSRS; (28)). For this study, only 14 items of the original 36 in the adult mSRS were selected, based on high intra-item reliability and after eliminating the questions not applicable for the juvenile age or were considered too subjective or anthropomorphic –e.g. wanders aimlessly from one activity to another; touches others in an unusual way; shows indiscriminate grooming-(**Table 2**). In addition, the items were rated on a 5-point Likert scale (1= not true 0%, 2= sometimes true 25%, 3= often true 50%, 4=almost always true 75%, and 5= always true 100%), instead of the 4-point scale used by Feczko and colleagues (28) to provide higher resolution in the ratings. For each juvenile rhesus macaque observed, the 14-item jmSRS instrument was completed once, after the fourth focal observation (**Table 2**).

**Table 2.**
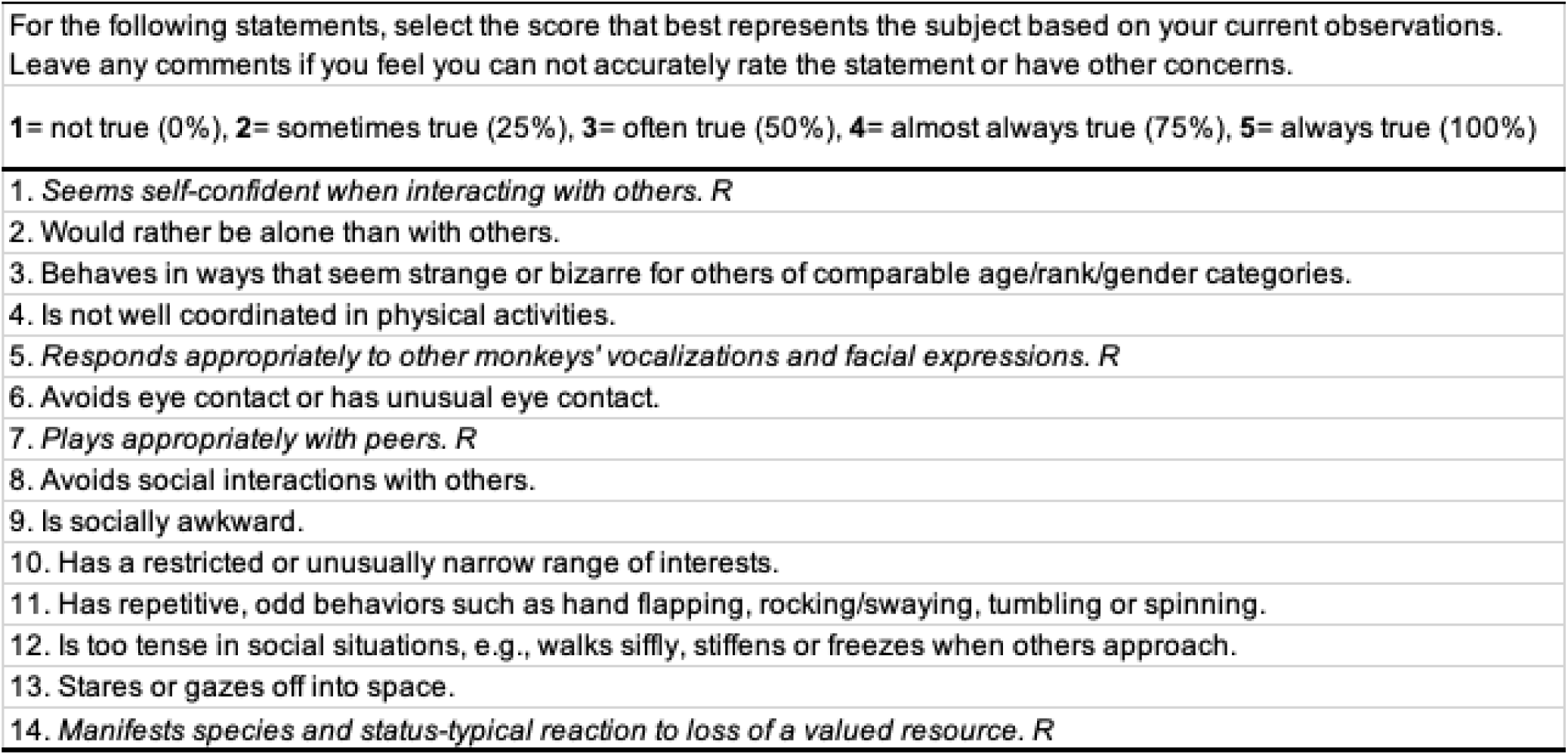
List of items included in the jmSRS and criteria for scoring based on the 1-5 Likert Rating Scales. The jmSRS was adapted from the adult mSRS (28). *R*: reverse-coded items (after data collection, the scoring of these items was reversed, so that higher scores meant greater social impairment for each item).

|  |
| --- |
| For the following statements, select the score that best represents the subject based on your current observations. Leave any comments if you feel you can not accurately rate the statement or have other concerns. |
| 1= not true (0%), 2= sometimes true (25%), 3= often true (50%), 4= almost always true (75%), 5= always true (100%) |
| 1. *Seems self-confident when interacting with others. R* |
| 2. Would rather be alone than with others. |
| 3. Behaves in ways that seem strange or bizarre for others of comparable age/rank/gender categories. |
| 4. Is not well coordinated in physical activities. |
| 5. *Responds appropriately to other monkeys' vocalizations and facial expressions. R* |
| 6. Avoids eye contact or has unusual eye contact. |
| 7. *Plays appropriately with peers. R* |
| 8. Avoids social interactions with others. |
| 9. Is socially awkward. |
| 10. Has a restricted or unusually narrow range of interests. |
| 11. Has repetitive, odd behaviors such as hand flapping, rocking/swaying, tumbling or spinning. |
| 12. Is too tense in social situations, e.g., walks siffly, stiffens or freezes when others approach. |
| 13. Stares or gazes off into space. |
| 14. *Manifests species and status-typical reaction to loss of a valued resource. R* |

### Data Reduction and Analysis

JmSRS data reduction and analyses of distribution, demographics, internal reliability and validity followed previously published methods by our group to develop and validate an instrument for global maternal quality rating in rhesus monkeys (35).

#### Descriptive Statistics of the jmSRS

Prior to analysis of the jmSRS data distribution, demographics, internal reliability and validity, four items were reverse-coded to match the interpretation of higher scores meaning greater social impairment by the rest of the items: (1) “Seems self-confident when interacting with others”, (5) “Responds appropriately to other monkeys’ vocalizations and facial expressions”, (7) “Plays appropriately with peers”, (14) “Manifests species and status-typical reaction to loss of a valued resource”. Total scores for each subject were calculated as the sum of all items, with possible scores ranging between 14 and 70 (based on 14 items with a 1-5 Likert scale score/each). The individual distribution of the total jmSRS scores were plotted and the Kolmogorov-Smirnov test was used to analyze the shape and distribution of the total scores.

#### Analyses of jmSRS Internal Reliability

Exploratory factor analysis (EFA) was performed using IBM SPSS Statistics 25 to determine the overall factor structure of the jmSRS 14 items and for data reduction. “Principal axis factoring” was used as the extraction method. The “latent root criterion” was initially used to establish the number of factors. It resulted in four factors; however, because there was a high number of cross-loadings, and only two items loaded on the fourth factor with a high factor weight (>0.3), we had to apply instead a criterion based on the “percentage of variance” to determine the number of factors to keep for analyses. Thus, we considered the solution that included just the first three EFA factors, which accounted for 60 percent of the total variance, as acceptable (see further details below, in Results). Promax rotation method was used to achieve simpler and theoretically more meaningful factor solutions. Factor scores were then calculated for each subject individually, using the factor score method.

After the factor structure of the jmSRS was determined using EFA, the internal reliability of each factor was tested using Cronbach’s α analysis. The internal consistency of a factor was considered high if α ≥0.7, moderate if 0.5 ≤ α < 0.69, or unacceptable if α < 0.5.

#### Analysis of jmSRS validity

In order to assess the convergent construct and discriminant validity of the jmSRS, we examined the associations between each jmSRS EFA factor score and the individual social and non-social behaviors collected during the two hours of focal observations with the ethogram. For that, the frequency or the duration of each behavior was first summed per subject across the two hours of behavioral observations, and then frequency rate or duration proportion per hour was calculated for each behavior. Then, to test the validity of the jmSRS, Spearman correlation coefficients were calculated to examine the associations between the jmSRS factor scores and frequency rates or proportion of time spent on behaviors collected during the focal behavioral observations. Correlation coefficients were calculated between behaviors in **Table 1** (except sitting alone and stereotypic behaviors, which were excluded due to low occurrence [<15%]) and the jmSRS factor scores. Significance level was set at p < 0.05.

#### Analyses of Sex and Social Rank Differences

Multivariate ANOVA was used to examine the effects of SEX (male, female) and social RANK (high, medium, low) factors on the jmSRS factor scores. A parallel multivariate ANOVA was performed on behavioral data collected with our ethogram to confirm expected SEX and Social RANK differences in this species. Before the ANOVAs, the Kolmogorov-Smirnov test was performed to examine the normal distribution of both jmSRS factor scores and behavioral frequency and duration rates; data were log-transformed when normality failed (p<0.05). Post-hoc pairwise comparisons were performed with Bonferroni adjustments, when main or interaction effects were detected. Significance level was set at p<0.05.

## Results

### JmSRS Total Scores

First, we calculated the sum of scores (total scores) for each animal. Scores ranged between 14 and 48, with a mean of 19.22 and a median of 18. The Kolmogorov-Smirnov analysis revealed that the total scores were not normally distributed (KS=0.186, p=1.5×10^−8^), and that the distribution was positively skewed (skew=1.868, kurtosis=0.759), indicating few animals (6.5%) had very high total scores (between 34-48; i.e. low in typical social behaviors and/or high in atypical behaviors). **Figure 1** shows the distribution of the jmSRS total scores.

**Fig 1.**
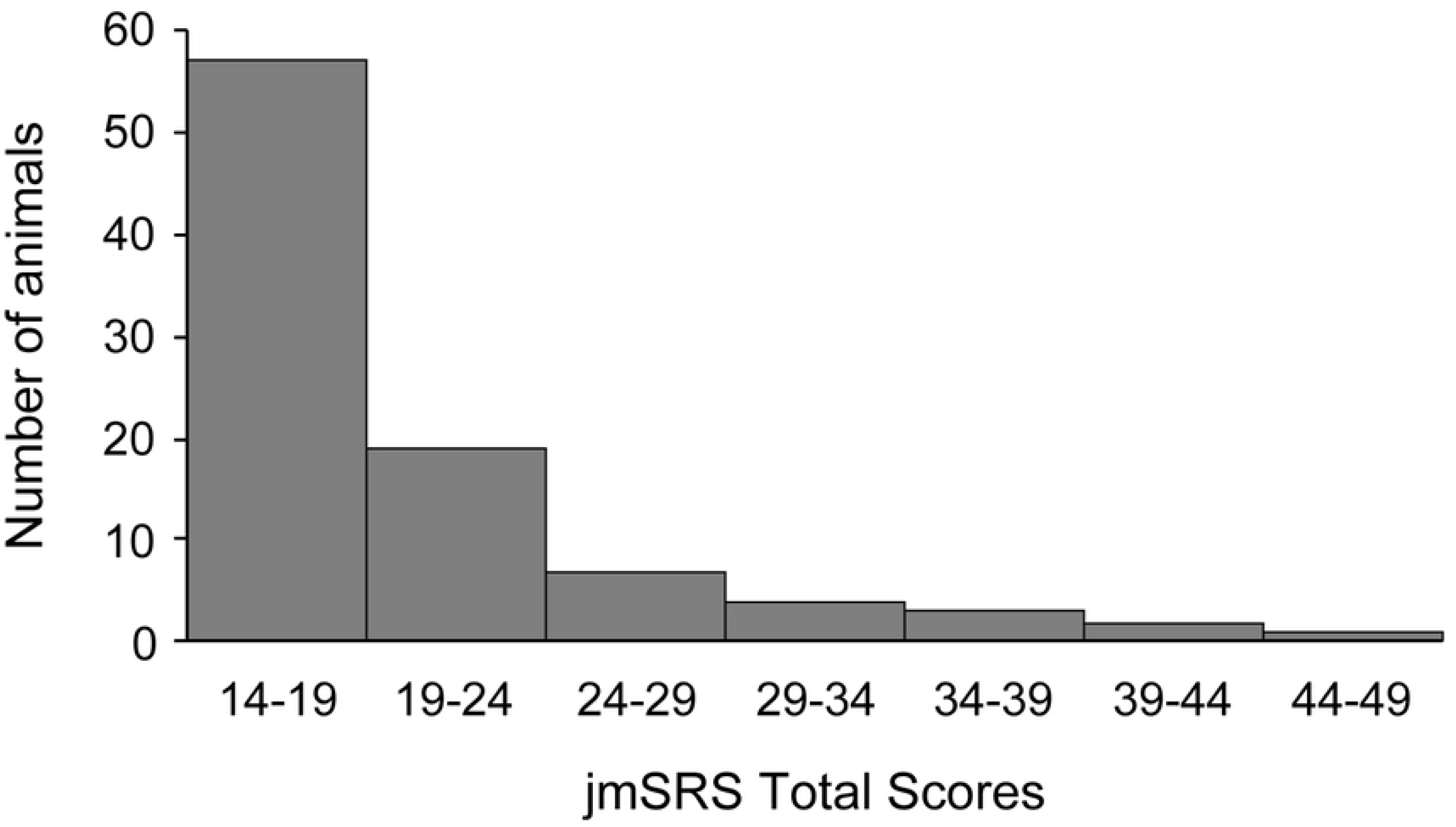
Histogram showing data frequency distribution of the jmSRS total scores in our juvenile macaque population. The jmSRS total score was calculated for each subject as the sum of scores for each jmSRS item. Each column represents the number of animals with the jmSRS total score indicated on the x-axis.

### JmSRS Exploratory Factor Analysis (EFA): Internal Reliability

The jmSRS EFA revealed a solution in which the first three factors explained approximately 60% of the variance in the data, accounting for 39.06%, 10.91% and 9.41%, respectively. In this selected 3-factor EFA solution (**Table 3**), jmSRS Factor #1 explained the majority of the variance -similarly to the human and chimpanzee SRS and the adult mSRS (26, 27, 36)-, and contained items that matched the first criterion domain of human ASD (deficits in social communication and social interactions; (35)) in the human SRS Factor 1, as well as items related to social avoidance and social anxiety in the mSRS and chimpanzee SRS Factor 1 (e.g. *“Avoids social interactions with others”*, “*Would rather be alone than with others”, “Socially awkward”*; *“Is too tense in social situations”*). Thus, this first factor seems to measure the same construct across species and macaque ages (28, 29, 36). JmSRS Factor #2 contained items related to impaired motor coordination, staring off into space and problems with social self-confidence, while items in Factor #3 matched the second diagnostic criteria of ASD (repetitive/stereotypic motor patterns; 35) and bizarre behaviors. Each item in the jmSRS had a high factor loading (>0.3) on at least one factor (**Table 3**), and only Item 8 (*“Avoids social interactions with others”*) showed significant loadings on more than one factor (cross-loading on Factors #1 and #3), supporting the choice of this EFA 3-factor solution as the best underlying relationship structure between items; this means more segregation of items into different behavioral domains in the jmSRS than in the “unitary factor solution” of the human, chimpanzee and adult mSRS. Item 8 was the only one with a significant negative factor loading (on Factor #3), suggesting that animals that show strange/repetitive behaviors do not necessarily avoid social interactions.

**Table 3.** Exploratory factor analysis (EFA) of the 14 jmSRS items revealed a 3-factor solution that explained 60% of the variance in the data with high factor loadings (>0.3) of items. Gray numbers indicate items with low factor loadings (<0.3). *: Significant cross-loading on Factors #1 and #3.

|  | jmSRS Factor |  |  |
| --- | --- | --- | --- |
|  | #1 | #2 | #3 |
| **Item 2.** Would rather be alone than with others. | 0.81 | 0.122 | -0.121 |
| **Item 5.** Responds appropriately to other monkeys' vocalizations and facial expressions. | 0.426 | -0.15 | 0.258 |
| **Item 6.** Avoids eye contact or has unusual eye contact. | 0.58 | -0.2 | 0.291 |
| **Item 7.** Plays appropriately with peers. | 0.63 | 0.05 | 0.092 |
| **Item 8.** Avoids social interactions with others. | 0.91 | 0.112 | -0.331* |
| **Item 9.** Is socially awkward. | 0.751 | -0.11 | 0.104 |
| **Item 10.** Has a restricted or unusually narrow range of interests. | 0.646 | 0.069 | 0.12 |
| **Item 12.** Is too tense in social situations, e.g., walks siffly, stiffens or freezes when others approach. | 0.732 | -0.005 | 0.05 |
| **Item 14.** Manifests species and status-typical reaction to loss of a valued resource. | 0.417 | -0.126 | 0.045 |
| **Item 1.** Seems self-confident when interacting with others. | 0.279 | 0.521 | 0.009 |
| **Item 4.** Is not well coordinated in physical activities. | -0.272 | 0.8 | 0.23 |
| **Item 13.** Stares or gazes off into space. | 0.299 | 0.303 | 0.159 |
| **Item 3.** Behaves in ways that seem strange or bizarre for others of comparable age/rank/gender categories. | 0.249 | 0.156 | 0.726 |
| **Item 11.** Has repetitive, odd behaviors such as hand flapping, rocking/swaying, tumbling or spinning. | -0.084 | 0.182 | 0.426 |

Kolmogorov-Smirnov tests showed that the jmSRS EFA factor scores were not normally distributed (jmSRS #1: KS=0.213, p=2.36×10^−11^; jmSRS #2: KS=0.194, p=2.45×10^−9^; jmSRS #3: KS=0.225, p=1.01×10^−12^), and that the distribution of each factor score was positively skewed (jmSRS #1: skew=1.886, kurtosis=3.534; jmSRS #2: skew=2.637, kurtosis=9.187; jmSRS #3: skew=2.190, kurtosis=5.394). **Figure 2** shows the distribution of the jmSRS factor scores and the individual data.

**Fig 2.**
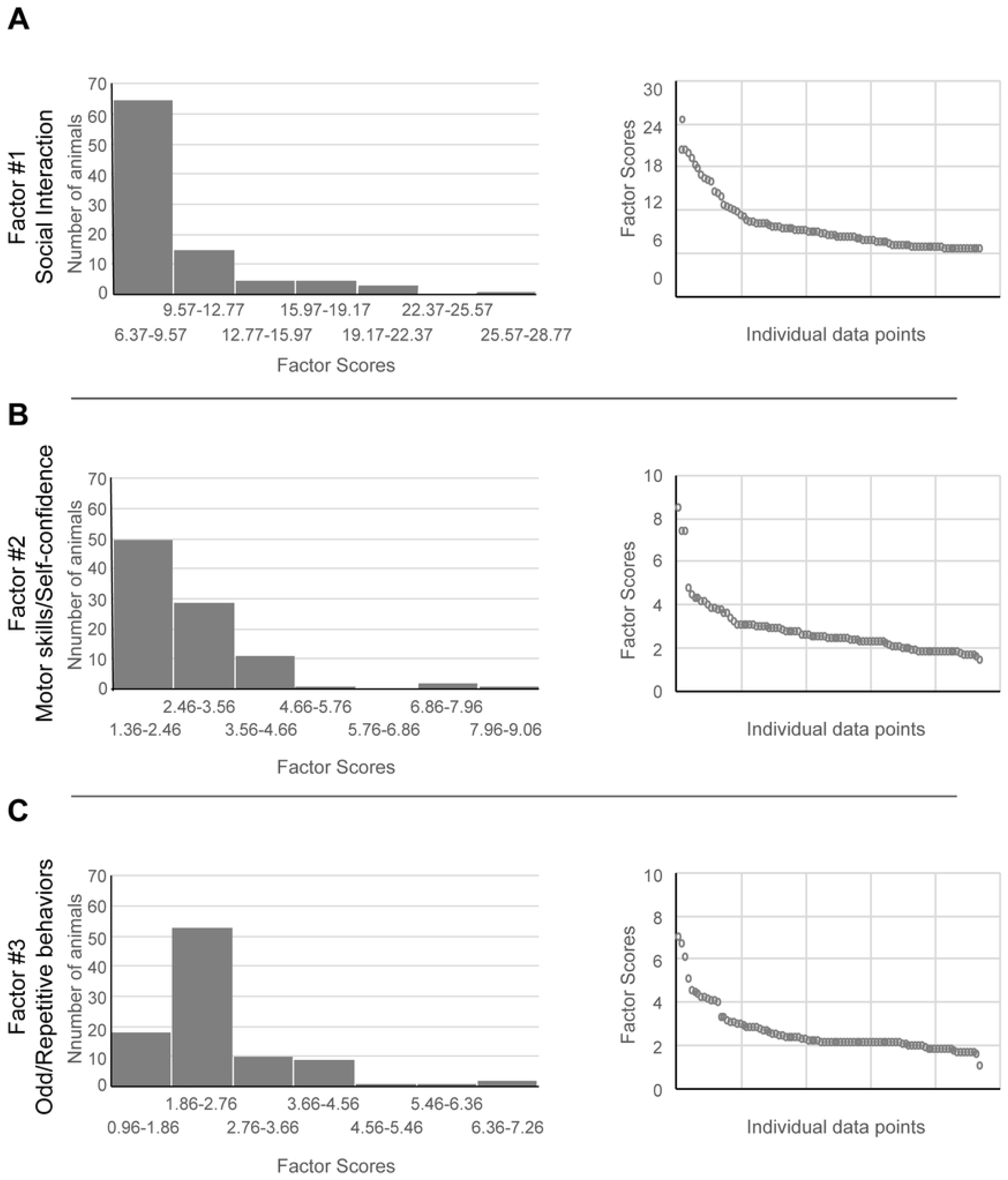
JmSRS EFA factors scores distributions. **Left**: Histograms showing data frequency distribution of jmSRS factors 1, 2 and 3’s scores (calculated by multiplying the item score by the factor loading per subject) in our juvenile macaque population. **Right**: Individual Factor scores, each circle represents a subject (n = 93). **A)** jmSRS Factor #1 (Social interaction), **B)** jmSRS Factor #2 (Motor coordination/Self-confidence), **C)** jmSRS Factor #3 (Odd/Repetitive behaviors).

Nine out of 14 items showed high factor loading on Factor #1, which represented the first criterion domain of ASD (social communication and interest), based on the DSM-V (3). Items loading on Factor #2 reflect self-confidence and motor coordination of the subject (“Seems self-confident when interacting with others”, “Is not well coordinated in physical activities”). Items loading on Factor #3 correlated to the second criterion domain of ASD as defined in the DSM-V (restrictive, repetitive behavior and interest, (3)). The factor score distributions along all three factors were positively skewed; that is, most of the subjects had low scores -indicating typical social responsiveness-, but for each factor several subjects in our population showed high scores, suggesting that each factor was able to detect outliers showing low typical and/or high atypical social behaviors (see **Figs. 2A, 2B, 2C**).

To test the internal (inter-item) consistency or reliability within each jmSRS factor, Cronbach α values were calculated. The analysis revealed good reliability for Factor #1, based on high Cronbach α (0.876), moderate reliability for Factor #2 (0.508) and low/unacceptable reliability for Factor #3 (0.436) – although, when the cross-loading item from Factor #3 is removed, the Cronbach’s α for Factor #3 increases to moderate levels (0.521).

Overall, the results of the EFA with a good 3-factor solution that explains approximately 60% of the variance and items with high factor loadings for each factor, and the high to acceptable Cronbach’s α suggest appropriate internal reliability of the three factors created from the 14-item jmSRS questionnaire.

### JmSRS Validity Analysis

Results of the correlation analysis between the jmSRS factors scores and the observed behaviors are showed in **Table 4**. JmSRS Factor #1 -which consists of items measuring impairments in social interactions and communication-showed significant positive correlations with *anxiety* (rho=0.276, p=0.008), *solitary play* behavior (rho=0.209, p=0.045), *following* behavior (rho=0.209, p=0.044), and *groom soliciting* (rho=0.218, p=0.035), whereas it was negatively correlated with *grooming* behavior (rho=-0.219, p=0.035). JmSRS Factor #2 -which represents items related to motor coordination, confidence, and staring off (which has been reported in children with ASD)-was positively correlated with *anxiety* (rho=0.323, p=0.002) and *groom soliciting* (rho=0.280, p=0.006), whereas it was negatively correlated with *aggressive behavior* (rho=-0.231, p<0.05). JmSRS Factor #3 -which represents items related to repetitive, odd behaviors and interests-was positively correlated with *anxiety* (rho=0.314, p=0.002), *groom soliciting* (rho=0.457, p=0.000), *solitary play* behavior (rho=0.320, p=0.002), *proximity* behavior (rho=0.280, p<0.007), *eye-gaze* behavior (rho=0.258, p=0.013), *breaking proximity* (rho=0.256, p=0.013), *following* behavior (rho=0.249, p=0.016) and *display* behavior (rho=0.330, p=0.001). These correlations suggest that impairments in social responsiveness in juveniles (i.e. high scores in the jmSRS Factors #1, #2 and #3) are associated with higher rates of anxiety, solitary play, and also breaking proximity with other animals. The significant correlations with social behaviors collected with the ethogram demonstrate the convergent/construct validity of the jmSRS instrument, and will be interpreted in more detail below, in the Discussion.

**Table 4.** Correlations between the jmSRS Factors (#1, #2, #3) and the behavior frequencies and durations collected with the Ethogram. **A)** Correlation matrix, showing significant correlations between behaviors and the jmSRS Factors (#1, #2, #3) in red font (p<.05); orange font: .05<p<.1; Spearman rho correlation coefficients. **B)** Examples of regression plots representing individual correlation data; the triangle, square and circle symbols represent the same 3 outliers in each graph.

| **A** |  |  | jmSRS Factors |  |  |
| --- | --- | --- | --- | --- | --- |
|  |  |  | FACTOR #1 | FACTOR #2 | FACTOR #3 |
| O<br>b<br>s<br>e<br>r<br>v<br>e<br>d<br>B<br>e<br>h<br>a<br>v<br>i<br>o<br>r<br>s | Groom soliciting | Spearman's rho | 0.218 | 0.28 | 0.457 |
|  |  | p value | 0.035 | 0.006 | 0.000 |
|  | Eye-gaze | Spearman's rho | 0.151 | 0.193 | 0.258 |
|  |  | p value | 0.147 | 0.064 | 0.013 |
|  | Grooming | Spearman's rho | -0.219 | -0.085 | 0.120 |
|  |  | p value | 0.035 | 0.417 | 0.250 |
|  | Proximity | Spearman's rho | -0.070 | -0.112 | 0.280 |
|  |  | p value | 0.503 | 0.283 | 0.007 |
|  | Solitary play | Spearman's rho | 0.209 | 0.166 | 0.320 |
|  |  | p value | 0.045 | 0.112 | 0.002 |
|  | Agression | Spearman's rho | -0.069 | -0.231 | 0.083 |
|  |  | p value | 0.510 | 0.026 | 0.429 |
|  | Display | Spearman's rho | 0.006 | 0.088 | 0.330 |
|  |  | p value | 0.954 | 0.402 | 0.001 |
|  | Follow | Spearman's rho | 0.209 | 0.093 | 0.249 |
|  |  | p value | 0.044 | 0.374 | 0.016 |
|  | Leave behind | Spearman's rho | 0.166 | 0.115 | 0.256 |
|  |  | p value | 0.111 | 0.273 | 0.013 |
|  | Anxiety | Spearman's rho | 0.276 | 0.323 | 0.314 |
|  |  | p value | 0.008 | 0.002 | 0.002 |

### Sex and Social Rank Differences

Based on the results of the Kolmogorov-Smirnov test described above, showing that the jmSRS EFA factor scores were not normally distributed (jmSRS #1: KS=0.213, p=2.36×10^−11^; jmSRS #2: KS=0.194, p=2.45×10^−9^; jmSRS #3: KS=0.225, p=1.01×10^−12^), multivariate ANOVA was run in log-transformed data. Results of the ANOVA showed no main SEX (jmSRS #1: F(1,85)=2.209, p=0.14; jmSRS #2: F(1,85)=0.437, p=0.51; jmSRS #3: F(1,85)=0.255, p=0.62) or RANK effects (jmSRS #1: F(2,85)=0.348, p=0.71; jmSRS #2: F(2,85)=1.152, p=0.32; jmSRS #3: F(2,85)=0.833, p=0.44) or SEX x RANK interaction effects (jmSRS #1: F(2,85)=0.315, p=0.73; jmSRS #2: F(2,85)=0.332, p=0.72; jmSRS #3: F(2,85)=0.109, p=0.90).

In contrast, a parallel multivariate ANOVA performed on behavioral data collected with our ethogram confirmed the expected SEX and Social RANK differences in this species. Because Kolmogorov-Smirnov test revealed non-normal distribution of behavioral data, and log-transformation could not be performed due to some behavioral scores being “0”, we used the Greenhouse-Geisser-corrected results. The ANOVA revealed significant SEX differences for the following behaviors: social play (F(1,84)=9.172, p=0.003, males>females), sexual F(1,84)=32.264, p=1.88×10^−7^, males>females), submissive behavior (F(1,84)=6.53, p=0.012, males<females) and fear grimace (F(1,84)=8.777, p=0.004, males<females), and a trend for breaking proximity (F(1,84)=3.37, p=0.07, males<females). We also found significant RANK effects for the following behaviors: aggression (F(2,84)=5.376, p=0.006, high>middle>low rank animals) and submissive behavior (F(2,84)=15.548, p=2×10^−6^, low>middle> high rank animals). Significant SEX x RANK interaction was observed for proximity behavior (F(2,84)=5.136, p=0.008), with more time spent in proximity for high ranking males than females, whereas the opposite was observed in middle ranking animals.

## Discussion

The goal of this study was to adapt and validate the adult mSRS to assess social and non-social behaviors in juvenile macaques of equivalent ages to 4-6 year old human children. We termed this downward extension of the instrument the “jmSRS”, composed of 14 items that measure global dimensions of typical and atypical social behaviors, as well as stereotypic and abnormal behaviors of relevance to ASD in juveniles. The EFA analysis identified 3 jmSRS factors with high levels of internal consistency/reliability and items with high factor loadings along the constructs identified in the human SRS as relevant to ASD: #1) impairments in social interactions and communication, #2) impairments in motor coordination, staring off, self-confidence, and #3) stereotypic/repetitive/odd behaviors and interests. The jmSRS identifies variability in social responsiveness in juvenile macaques and shows sensitivity to detect individuals with atypical social behaviors in both males and females, and across social ranks. Finally, the significant correlations between impairments in social responsiveness in the 3 jmSRS factors and impairments in social and motor behaviors and higher rates of anxiety detected with behavioral ethogram demonstrate the convergent construct validity of the jmSRS instrument. Our findings indicate that the jmSRS can be easily filled by trained primate researchers, complementing standard focal observations with social responsiveness data critical for future screening in macaque models of ASD.

The human SRS is a widely used rating scale to quantitatively measure the continuum of symptom severity in ASD in children (8). This test is usually used with other diagnostic tools to assess of symptom severity in ASD in children, but rarely in adults (11). The complex and neurodevelopmental nature of ASD has made it hard to study its biological roots in children. Thus, the study of NHP models including the rhesus monkey, a species that closely resembles humans in terms of socially complex behavior, brain anatomy/connectivity and development, is crucial to understand neurobiological mechanisms underlying ASD. Cross-species adaptations of the human SRS have been published in chimpanzees (29) and rhesus macaques (mSRS; (28)); however, both studies focused on adapting the SRS to adult populations. Given that the early diagnosis and assessment of ASD symptom severity is crucial for intervention in children and infants, this study adapted and validated the adult mSRS for juvenile male and female macaques of all social status/ranks at approximately 1.5 yrs, which is roughly equivalent to 4-6 yrs of age in children, with the goal of using this screening instrument to identify atypical social, motor and other behaviors of relevance for ASD in children among socially-housed juvenile macaques.

The 36 original items of the adult mSRS were evaluated and adapted for fit to the juvenile developmental period, and a 14-item jmSRS rating scale was created. Selection of items was based on high intra-item reliability, applicability to juveniles and non-subjective or anthropomorphic questions. Trained observers scored the social behavior of 93 juvenile rhesus macaques using the jmSRS, in parallel to traditional behavioral observation data collected using well-established ethograms (33, 34, 36). Analysis of the distribution of jmSRS total scores (sum of scores of all items per subject) revealed positively skewed, not normally distributed total scores, similarly to the adult mSRS (28) and the human SRS (37-39). This result indicates that very few animals (6.5%) received very high total jmSRS scores - between 34 and 48-(i.e. high scores in atypical behaviors); most juveniles showed high typical and low atypical social behaviors, which is consistent with the human SRS observations (37).

The exploratory factor analysis (EFA) revealed a good 3-factor solution that explained approximately 60% of the variance, had high factor loadings for items and high to acceptable internal consistency/reliability. Factor #1 represented items that matched the first major criterion domain of ASD (deficits in social communication and social interactions, based on the DSM-5 (3)), and accounted for the vast majority of the variance (39.06%), similarly to the human (34.97%), the chimpanzee (52.27%) and the adult mSRS (30.64%), whereas the remaining factors explained the minority of the variance in the jmSRS and the other studies (28, 29, 40). Factor #2 represented items related to impaired motor coordination –consistent with recent reports of impaired motor coordination, postural control and balance in children with ASD (41-43), staring off into space– also consistent with reports in children with ASD (44), and self-confidence when interacting with others. Factor #3 included items represented in the second diagnostic criteria of ASD (restricted, repetitive/stereotypic patterns of behavior, interests or activities (3)). This 3-factor structure recapitulates what earlier studies revealed in adult macaques and chimpanzees (the human SRS showed a 5-factor structure (40)), except for a key difference: although the jmSRS had two separate factors representing the two major ASD criterion domains (Factors #1 and #3), in the human, chimpanzee and the adult mSRS both criterion domains are included in the first factor (28, 29, 40). A cross-species comparison revealed that items loading strongly on Factor #1 in all species (independently of age) are related to social communication and interactions, such as “avoids eye-contact”, “avoids social interaction”, and “socially awkward”, which confirms that this factor reliably identifies similarities in social behavior across primate species (8, 28, 29). This observation highlights the translational value of the instrument for ASD in humans.

The distribution of the jmSRS factor scores were also positively skewed, with the vast majority of the subjects (Factor#1: 80%, Factor#2: 82%, Factor#3: 73%) showing low scores, indicating species-typical and age-appropriate social behavior. Yet, few subjects scored high on Factor#1 (4.3%), Factors #2 and #3 (3.2%) of the jmSRS, supporting the sensitivity of this jmSRS instrument to identify outliers based on their atypical social, motor and stereotypic/odd behavior. The lack of sex or social rank effects on the jmSRS factor scores, despite confirmation of expected sex and social rank differences in behavioral data collected with our ethogram (34, 36) was surprising, particularly given the association between higher scores and lower rank in the adult mSRS (28), the chimpanzee SRS (29), and the male:female higher ratio of children with ASD deficits (2, 8). Future studies with larger samples sizes need to further examine these relationships.

The significant correlations between impairments in social responsiveness, motor coordination and stereotypic/repetitive behavior in the jmSRS factors and low sociability, high anxiety data collected during focal behavioral observations of each subject using well-established ethograms demonstrate the convergent construct validity of the jmSRS instrument. Subjects scoring higher on the jmSRS Factor #1 (i.e. showing deficits in social communication and interactions) were more anxious, played alone for longer periods of time, and groomed other animals less. Under normal circumstances, grooming behavior in this species emerges around five months of age and increases with age (18). First, infants practice grooming behavior on their mother, which later gets generalized to other family members (aunts, cousins), and then to peers and other animals in the subject’s environment (18). Playing behavior in macaques exhibits a similar developmental increase, beginning around 7-12 weeks of age followed by a rapid increase in the frequency until it peaks near the end of the first year, which coincides with the age at which our subjects were observed (18). Thus, decreased amounts of grooming and increased amounts of non-social (solitary) play suggest lower sociability, which confirms the construct validity and sensitivity of the first jmSRS factor to identify deficits in social communication and interactions. Given that ASD is defined by impairments in social communication and interactions, in parallel to restricted interests, repetitive patterns of behavior and interests (3, 4), our findings suggest that the jmSRS is a promising translational tool for studies in developing macaque models of children ASD.

Subjects with higher scores on Factor #2 (not well-coordinated, staring off into space and less confident animals) were also more anxious and less likely to be aggressive. In macaques, aggressive behavior first appears around 30 weeks of age, mostly towards siblings and less dominant peers, as a playful way of practice (18). Interestingly, macaques with specific early experiences, particularly insecure early attachment with their mothers, tend to exhibit aggressive behavior later (45). In juvenile-age macaques, peer-related activities – such as aggression – continue to be present at high rates until puberty (46), which suggests that animals exhibiting lower amounts of aggressive behavior might be more immature or have poorer social skills. Impaired motor coordination, postural control and balance has been reported in children with ASD (41-43), as well as staring off into space (47, 48).

Finally, subjects exhibiting more repetitive, atypical behaviors (Factor #3) were also more anxious (based on our measures of anxiety-like behaviors –scratching/yawning/body shake/self-directed behaviors) and played alone for longer periods, but they were also seeking others’ attention/proximity more (more eye-gazing, following, proximity and groom-soliciting behaviors). The associations between high scores in the 3 jmSRS factors and elevated anxiety is very interesting, as a recent meta-analysis has concluded that almost 40% of children and adolescents with ASD exhibit symptoms of anxiety (4). Both “deficits in social interaction and communication” (49, 50) and “restrictive and repetitive behaviors” (51-53) –which are the two major criteria for ASD diagnoses (3)– have been associated with increased levels of anxiety. A recent study that analyzed the association between social responsiveness using the human SRS and anxiety in 150 pre-adolescents and adolescents with ASD, also showed strong correlations between the SRS total score and anxiety, and between the “Social communication” and “Autistic mannerisms” subscales of the SRS and anxiety (54), which is consistent with the associations we report here, in our juvenile macaque population.

The goal of this study was to adapt and test the validity of the jmSRS, a downward extension of the adult mSRS (28) to juvenile-age macaques, as a potential sensitive instrument that provides measures of atypical social, motor and stereotypic behaviors, similarly to the human SRS used for ASD diagnosis in children (8, 40). The ultimate goal is to develop and apply this translational tool to identify social deficits of relevance for ASD in socially-housed macaque NHP models. Developing translational NHP models of ASD-related social alterations is critical, and such models must show similar patterns and complexity of brain and social development to humans. As detailed above, the 14-item jmSRS developed in this study showed several similarities but also differences compared to the SRS used in human studies and validated in chimpanzee and adult macaque studies earlier. Although the first factor correlates with the major diagnostic criteria of ASD in both human and the adult NHP studies, the present study revealed two separate factors (Factor #1: social interactions and #3: repetitive/odd behaviors) representing the two major ASD criteria domains in the jmSRS. Correlations between jmSRS Factor #1 scores and behavioral frequency and duration data collected with our ethogram showed that deficits in social communication and interactions was positively correlated with “anxiety” and “solitary play behavior”, but negatively correlated with “grooming”; Factor #2 was more strongly and positively correlated with “anxiety” and “groom soliciting behaviors”; and Factor # 3 -which measures odd/repetitive behaviors - was strongly and positively correlated with “anxiety”, “leave behind” and “solitary play behaviors”, showing similar associations than those reported for Factor #1. These correlations support that the jmSRS reliably measures atypical social behaviors present in ASD.

The study also has some limitations that need to be considered. First, we were not able to address the intra-rater reliability in the jmSRS scores. However, the experimental design involved expert macaque behavior coders that filled in the jmSRS questionnaire after completion of all 4 × 30 min focal behavioral observations based on our ethogram (so they were very familiar with juvenile macaque -and each subjects’-social behavior), and their inter-rater reliability for those focal observations was high (Cohen’s k >.80). Future studies need to address both the jmSRS inter- and intra-rater reliability across all coders. Additional studies are also needed to adapt the jmSRS instrument to measure typical and atypical behaviors in younger –infant-monkeys.

In summary, this study adapted a downward extension of the adult mSRS, and tested its external validity and sensitivity to identify atypical social, motor and stereotypic behaviors of relevance to ASD in socially-housed juvenile rhesus macaques. Our findings with the jmSRS instrument show high levels of internal reliability, sensitivity to detect individuals with atypical behaviors and convergent construct validity, confirmed by the associations between the jmSRS factors and the behavioral observation data. Based on the similarities found with the human SRS, the jmSRS is a promising translational tool for studies in developing macaque models of ASD in children.

## Acknowledgements

The authors want to thank Jennifer Whitley, Josh Bailey, Manuel Bautista and Jessica Johnson, members of the YNPRC Field Station Phenotyping group (PhenX: Identification of Unique Phenotypes at the YNPRC Breeding Colony), and Colony Management for collection of behavioral data; Dr. Rebecca Herman for developing the data extraction/error checking programs; and Drs. Hasse Walum and Melinda Higgins for statistical guidance.

## References

1. Reeve SA, Reeve KF, Townsend DB, Poulson CL. Establishing a generalized repertoire of helping behavior in children with autism. J Appl Behav Anal. 2007;40(1):123–36.

2. Baio J, Wiggins L, Christensen DL, Maenner MJ, Daniels J, Warren Z, et al. Prevalence of Autism Spectrum Disorder Among Children Aged 8 Years - Autism and Developmental Disabilities Monitoring Network, 11 Sites, United States, 2014. MMWR Surveill Summ. 2018;67(6):1–23.

3. 5th edition ed. Arlington, VA: American Psychiatric Publishing; 2013. Diagnostic and statistical manual of mental disorders (DSM-V).

4. van Steensel FJ, Bogels SM, Perrin S. Anxiety disorders in children and adolescents with autistic spectrum disorders: a meta-analysis. Clin Child Fam Psychol Rev. 2011;14(3):302–17.

5. Lai MC, Lombardo MV, Chakrabarti B, Ruigrok AN, Bullmore ET, Suckling J, et al. Neural self-representation in autistic women and association with ‘compensatory camouflaging’. Autism. 2019;23(5):1210–23.

6. Pickles A, Le Couteur A, Leadbitter K, Salomone E, Cole-Fletcher R, Tobin H, et al. Parent-mediated social communication therapy for young children with autism (PACT): long-term follow-up of a randomised controlled trial. Lancet. 2016;388(10059):2501–9.

7. Wetherby AM, Guthrie W, Woods J, Schatschneider C, Holland RD, Morgan L, et al. Parent-implemented social intervention for toddlers with autism: an RCT. Pediatrics. 2014;134(6):1084–93.

8. Constantino JN, Davis SA, Todd RD, Schindler MK, Gross MM, Brophy SL, et al. Validation of a brief quantitative measure of autistic traits: comparison of the social responsiveness scale with the autism diagnostic interview-revised. J Autism Dev Disord. 2003;33(4):427–33.

9. Kamp-Becker I, Albertowski K, Becker J, Ghahreman M, Langmann A, Mingebach T, et al. Diagnostic accuracy of the ADOS and ADOS-2 in clinical practice. Eur Child Adolesc Psychiatry. 2018;27(9):1193–207.

10. Mazefsky CA, Oswald DP. The discriminative ability and diagnostic utility of the ADOS-G, ADI-R, and GARS for children in a clinical setting. Autism. 2006;10(6):533–49.

11. Chan W, Smith LE, Hong J, Greenberg JS, Mailick MR. Validating the social responsiveness scale for adults with autism. Autism Res. 2017;10(10):1663–71.

12. Bowlby J. Attachment theory and its therapeutic implications. Adolesc Psychiatry. 1978;6:5–33.

13. Harlow HF, Zimmermann RR. Affectional responses in the infant monkey; orphaned baby monkeys develop a strong and persistent attachment to inanimate surrogate mothers. Science. 1959;130(3373):421–32.

14. Meyer JS, Novak MA, Bowman RE, Harlow HF. Behavioral and hormonal effects of attachment object separation in surrogate-peer-reared and mother-reared infant rhesus monkeys. Dev Psychobiol. 1975;8(5):425–35.

15. Chang SW, Barter JW, Ebitz RB, Watson KK, Platt ML. Inhaled oxytocin amplifies both vicarious reinforcement and self reinforcement in rhesus macaques (Macaca mulatta). Proc Natl Acad Sci U S A. 2012;109(3):959–64.

16. Chang SW, Winecoff AA, Platt ML. Vicarious reinforcement in rhesus macaques (macaca mulatta). Front Neurosci. 2011;5:27.

17. Ferrari PF, Paukner A, Ionica C, Suomi SJ. Reciprocal face-to-face communication between rhesus macaque mothers and their newborn infants. Curr Biol. 2009;19(20):1768–72.

18. Hinde RA, Spencer-Booth Y. The behaviour of socially living rhesus monkeys in their first two and a half years. Anim Behav. 1967;15(1):169–96.

19. Damon F, Meary D, Quinn PC, Lee K, Simpson EA, Paukner A, et al. Preference for facial averageness: Evidence for a common mechanism in human and macaque infants. Sci Rep. 2017;7:46303.

20. Machado CJ, Bachevalier J. Non-human primate models of childhood psychopathology: the promise and the limitations. J Child Psychol Psychiatry. 2003;44(1):64–87.

21. Parr LA. The evolution of face processing in primates. Philosophical transactions of the Royal Society of London Series B, Biological sciences. 2011;366(1571):1764–77.

22. Passingham R. How good is the macaque monkey model of the human brain? Current opinion in neurobiology. 2009;19(1):6–11.

23. Preuss TM. Taking the measure of diversity: comparative alternatives to the model-animal paradigm in cortical neuroscience. Brain, behavior and evolution. 2000;55(6):287–99.

24. Suomi SJ. Risk, resilience, and gene x environment interactions in rhesus monkeys. Ann N Y Acad Sci. 2006;1094:52–62.

25. Bauman MD, Schumann CM. Advances in nonhuman primate models of autism: Integrating neuroscience and behavior. Exp Neurol. 2018;299(Pt A):252–65.

26. Sanchez MM, Ladd CO, Plotsky PM. Early adverse experience as a developmental risk factor for later psychopathology: evidence from rodent and primate models. Dev Psychopathol. 2001;13(3):419–49.

27. Zhu Y, Sousa AMM, Gao T, Skarica M, Li M, Santpere G, et al. Spatiotemporal transcriptomic divergence across human and macaque brain development. Science. 2018;362(6420).

28. Feczko EJ, Bliss-Moreau E, Walum H, Pruett JR, Jr., Parr LA. The Macaque Social Responsiveness Scale (mSRS): A Rapid Screening Tool for Assessing Variability in the Social Responsiveness of Rhesus Monkeys (Macaca mulatta). PLoS One. 2016;11(1):e0145956.

29. Marrus N, Faughn C, Shuman J, Petersen SE, Constantino JN, Povinelli DJ, et al. Initial description of a quantitative, cross-species (chimpanzee-human) social responsiveness measure. J Am Acad Child Adolesc Psychiatry. 2011;50(5):508–18.

30. Maestripieri D. Parenting styles of abusive mothers in group-living rhesus macaques. Anim Behav. 1998;55(1):1–11.

31. McCormack K, Sanchez MM, Bardi M, Maestripieri D. Maternal care patterns and behavioral development of rhesus macaque abused infants in the first 6 months of life. Dev Psychobiol. 2006;48(7):537–50.

32. Council NR. Guide for the Care and Use of Laboratory Animals. Washington, DC: National Academies Press; 2011. Report No.: ISBN-13: 978-0-309-15400-0.

33. Altmann SA. A Field Study of the Sociobiology of Rhesus Monkeys, Macaca Mulatta. Annals of the New York Academy of Sciences. 1962;102:338–435.

34. Herman RA, Measday MA, Wallen K. Sex differences in interest in infants in juvenile rhesus monkeys: relationship to prenatal androgen. Horm Behav. 2003;43(5):573–83.

35. McCormack K, Howell BR, Guzman D, Villongco C, Pears K, Kim H, et al. The development of an instrument to measure global dimensions of maternal care in rhesus macaques (Macaca mulatta). Am J Primatol. 2015;77(1):20–33.

36. Raper J, Stephens SB, Sanchez M, Bachevalier J, Wallen K. Neonatal amygdala lesions alter mother-infant interactions in rhesus monkeys living in a species-typical social environment. Dev Psychobiol. 2014;56(8):1711–22.

37. Constantino JN, Todd RD. Autistic traits in the general population: a twin study. Arch Gen Psychiatry. 2003;60(5):524–30.

38. Cheon KA, Park JI, Koh YJ, Song J, Hong HJ, Kim YK, et al. The social responsiveness scale in relation to DSM IV and DSM5 ASD in Korean children. Autism Res. 2016;9(9):970–80.

39. Constantino JN. The quantitative nature of autistic social impairment. Pediatr Res. 2011;69(5 Pt 2):55R–62R.

40. Constantino JN, Gruber CP, Davis S, Hayes S, Passanante N, Przybeck T. The factor structure of autistic traits. J Child Psychol Psychiatry. 2004;45(4):719–26.

41. Bedford R, Pickles A, Lord C. Early gross motor skills predict the subsequent development of language in children with autism spectrum disorder. Autism Res. 2016;9(9):993–1001.

42. Buja A, Volfovsky N, Krieger AM, Lord C, Lash AE, Wigler M, et al. Damaging de novo mutations diminish motor skills in children on the autism spectrum. Proc Natl Acad Sci U S A. 2018;115(8):E1859–E66.

43. Fulceri F, Grossi E, Contaldo A, Narzisi A, Apicella F, Parrini I, et al. Motor Skills as Moderators of Core Symptoms in Autism Spectrum Disorders: Preliminary Data From an Exploratory Analysis With Artificial Neural Networks. Front Psychol. 2018;9:2683.

44. Hughes R, Poon WY, Harvey AS. Limited role for routine EEG in the assessment of staring in children with autism spectrum disorder. Arch Dis Child. 2015;100(1):30–3.

45. Suomi SJ. Aggression and social behaviour in rhesus monkeys. Novartis Found Symp. 2005;268:216–22; discussion 22-6, 42-53.

46. Suomi SJ. Mother-Infant Attachment, Peer Relationships, and the Development of Social Networks in Rhesus Monkeys. Human Development. 2005;48:67–79.

47. Keller R, Basta R, Salerno L, Elia M. Autism, epilepsy, and synaptopathies: a not rare association. Neurol Sci. 2017;38(8):1353–61.

48. Ko C, Kim N, Kim E, Song DH, Cheon KA. The effect of epilepsy on autistic symptom severity assessed by the social responsiveness scale in children with autism spectrum disorder. Behav Brain Funct. 2016;12(1):20.

49. Bellini S. Social Skill Deficits and Anxiety in High-Functioning Adolescents With Autism Spectrum Disorders. Focus on Autism and Other Developmental Disabilities. 2004;19(2):78–86.

50. Davis TE, 3rd, Moree BN, Dempsey T, Hess JA, Jenkins WS, Fodstad JC, et al. The effect of communication deficits on anxiety symptoms in infants and toddlers with autism spectrum disorders. Behav Ther. 2012;43(1):142–52.

51. Gotham K, Bishop SL, Hus V, Huerta M, Lund S, Buja A, et al. Exploring the relationship between anxiety and insistence on sameness in autism spectrum disorders. Autism Res. 2013;6(1):33–41.

52. Rodgers J, Riby DM, Janes E, Connolly B, McConachie H. Anxiety and repetitive behaviours in autism spectrum disorders and williams syndrome: a cross-syndrome comparison. J Autism Dev Disord. 2012;42(2):175–80.

53. Joyce C, Honey E, Leekam SR, Barrett SL, Rodgers J. Anxiety, Intolerance of Uncertainty and Restricted and Repetitive Behaviour: Insights Directly from Young People with ASD. J Autism Dev Disord. 2017;47(12):3789–802.

54. Bitsika V, Sharpley CF. The association between parents’ ratings of ASD symptoms and anxiety in a sample of high-functioning boys and adolescents with Autism Spectrum Disorder. Res Dev Disabil. 2017;63:38–45.

